# Cellular Dynamics of Skeletal Muscle Regeneration

**DOI:** 10.1101/2023.05.02.538744

**Authors:** Brittany C. Collins, Jacob B. Shapiro, Mya M. Scheib, Robert V. Musci, Mayank Verma, Gabrielle Kardon

## Abstract

The function of many organs, including skeletal muscle, depends on its three-dimensional structure. Muscle regeneration therefore requires not only reestablishment of myofibers, but restoration of tissue architecture. Resident muscle stem cells (SCs) are essential for regeneration, but how SCs regenerate muscle architecture is largely unknown. We address this problem using genetic labeling of SCs and whole mount imaging to reconstruct in three dimensions muscle regeneration. Unexpectedly, we found that the residual basement membrane of necrotic myofibers is critical for promoting fusion and orienting regenerated myofibers and myofibers form via two distinct phases of fusion. Furthermore, the centralized myonuclei characteristic of regenerated myofibers are associated with myofibrillogenesis and permanently mark regenerated myofibers. Finally, we elucidate a new cellular mechanism for the formation of branched myofibers, a pathology characteristic of diseased muscle. We provide a new synthesis of the cellular events of regeneration and show these differ from those used during development.

## INTRODUCTION

Stem cells are critical for regeneration of a variety of tissues. These cells are characterized by their ability to self-renew and to differentiate and regenerate tissue ^1^. To repair three-dimensionally complex tissues stem cells must not only differentiate, but they also need to appropriately restore tissue architecture. While the molecular and cellular processes regulating stem cell self-renewal and differentiation have been intensely studied, how stem cells regenerate tissue architecture is largely unknown. Here we examine in vivo how stem cells regenerate the architecture of a three-dimensionally complex tissue, skeletal muscle.

Adult vertebrate skeletal muscle is composed of multinucleate post-mitotic myofibers. In response to injury or damage, muscle has a remarkable capacity for regeneration that is mediated by a tissue resident population of dedicated muscle stem cells (SCs), originally termed satellite cells ^2–5^. Genetic lineage and ablation studies in mice have definitively established that SCs are the stem cells necessary and sufficient for muscle regeneration ^6–8^. In uninjured muscle, quiescent SCs reside in a “satellite” niche between the sarcolemma and the basement membrane (BM) of the myofiber ^5^. The BM is a sheath of extracellular matrices, notably laminin, that coats the basal aspect of myofibers ^9^. Upon muscle injury, SCs activate and re-enter the cell cycle. Asymmetric cell division of activated SCs is thought to establish the subsequent fate of SCs to either self-renew or differentiate into myofibers ^10–12^. SCs destined to regenerate myofibers become committed myoblasts, differentiate into myocytes, and fuse to reestablish myofibers ^13^.

SCs not only undergo myogenesis, but they correctly regenerate the three-dimensional muscle architecture. Each anatomical muscle has a unique organization and orientation of myofibers, which span from origin to distal tendons. After most forms of muscle injury, regeneration yields muscles with correctly oriented and aligned myofibers ^14, 15^. Histological and electron micrograph studies have suggested that the residual BM of necrotic myofibers is important for ensuring proper alignment of regenerated myofibers ^14, 16–19^, but these techniques lack the three-dimensional view required to reveal the complete regenerative process at the tissue level. Also essential for optimal muscle function is the regeneration of linear, as opposed to branched, myofibers—as the latter are associated with impaired contractile activity and more susceptible to damage ^20–23^. Branched myofibers are characteristic of dystrophic muscle ^23–30^ and are found with a higher incidence after regeneration ^28, 31^ and with age ^28^. However, the mechanisms that ensure production of linear versus branched regenerated myofibers are still unclear. Finally, it has long been recognized that the cytoarchitecture of regenerated myofibers is distinct from uninjured myofibers; regenerated myofibers have chains of centralized myonuclei, while uninjured myofibers have regularly spaced peripheral nuclei ^32–35^. How these centralized myonuclei arise during regeneration and whether (and how) these central nuclei move peripherally as regenerated myofibers mature are still outstanding questions ^33, 34, 36–38^.

The cellular events of regeneration are often suggested to closely mirror development ^39, 40^. The ultimate result of muscle development and regeneration is the creation of functional skeletal muscles. For the most part, the structure and function of developed and regenerated muscles look similar. However, there are some key differences, for instance the organization of myonuclei ^32, 33–35^. These morphological differences suggests that myogenesis may differ between regeneration and development.

In this study, using genetic labeling of SCs and imaging and reconstruction of muscle in whole mount, we determine how after myofiber destruction SCs regenerate the three-dimensional architecture of skeletal muscle. We provide a new synthesis of the cellular events leading to regeneration and reveal key differences between development and regeneration of skeletal muscle.

## RESULTS

### Three-dimensional visualization and quantification of myogenic cells during regeneration

To investigate SCs and their progeny during muscle regeneration in vivo, we optimized a previously published whole-mount tissue clearing technique ^41^ that allows for the three-dimensional visualization of the cellular processes of regeneration. We used a *Pax7^CreERT^*^2^ allele that specifically and efficiently induces Cre-mediated recombination in all SCs following tamoxifen (TAM) delivery ^7^. These mice were crossed to the *Rosa^mTmG^* reporter, which in the absence of Cre ubiquitously expresses membrane-bound Tomato (TOM), but in the presence of Cre, cells and their progeny express membrane-bound GFP ^42^. *Pax7^CreERT^*^2^*^/+^;Rosa^mTmG/+^* mice were used in all subsequent studies unless otherwise stated. We focused on regeneration of the tibialis anterior (TA) and extensor digitorum longus (EDL) muscles, whose myofibers are aligned parallel to one another and to the limb’s long axis. We genetically labeled SCs and their progeny with GFP via TAM, injured both TA and EDL muscles via intramuscular injection of BaCl2, and harvested muscles at various days post injury (DPI; Fig.1A, 2I, 3A). BaCl2 leads to acute damage of myofibers via cell membrane depolarization, hypercontraction, proteolysis, and membrane rupture ^43^. TA muscles were weighed and processed for fluorescence activated cell sorting (FACS) of SCs (gating strategy in Fig. S1), while EDL muscles were processed for whole-mount imaging.

**Figure 1.**
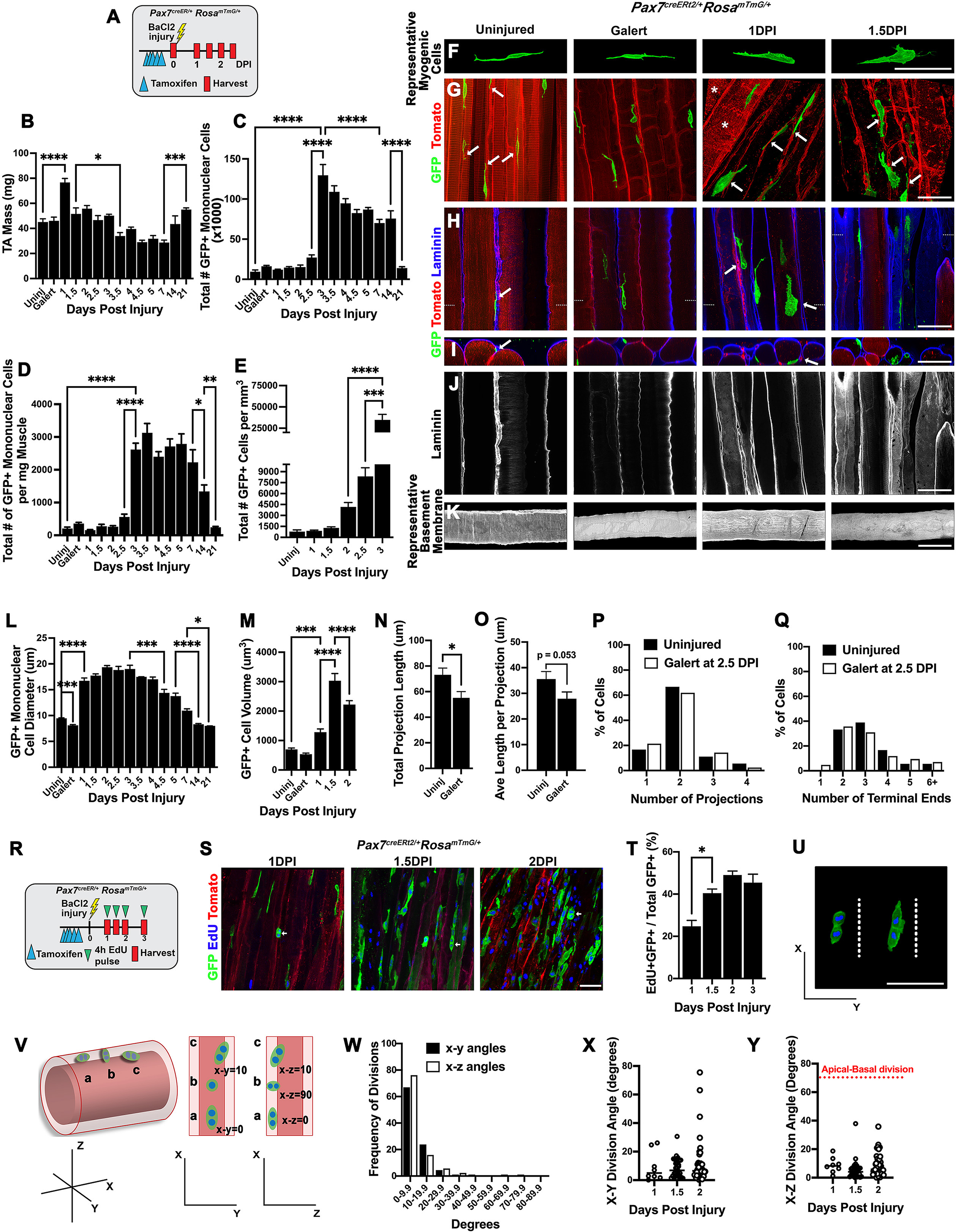
Quiescent, Galert, and activated SCs are distinct in size and shape and activated SCs proliferate via planar divisions within residual BM tubes. **(A)** Experimental design for genetic labeling of SCs, 0-2.5 DPI. **(B)** TA mass during regeneration (n = 3-5 mice/DPI). **(C-E)** Number of GFP+ mononuclear myogenic cells in TA by FACS (n = 3-5 mice/DPI) **(C-D)** and counts on wholemount images in EDL (n = 3-7 mice/DPI) **(E)**. **(F-K)** Representative wholemount images of uninjured, Galert (2.5 DPI contralateral uninjured EDL),1 DPI, and 1.5 DPI EDL (n= ≥ 3 mice/DPI). **(F)** Representative segmented SCs. **(G-H)** Superficial view of GFP+ SCs and TOM+ uninjured myofibers (Uninjured and Galert columns), TOM+ degrading myofibers (*’s at 1 DPI), and TOM+ capillaries (all columns). **(H)** Laminin+ BM outlines uninjured myofibers, residual BM of injured myofibers, and surrounds capillaries. Dotted white lines indicate level of cross-section shown in **(I)**. White arrows in **(G-I)** show GFP+ SCs in close contact with TOM+ capillaries. **(J)** Superficial view with only laminin of scans in **(H)**. **(K)** Representative segmented laminin+ BM of uninjured myofibers or residual BM of injured myofibers. **(L)** Cell diameter of GFP+ mononuclear myogenic cells measured via FACS of TA (n= 2-5 mice/DPI). **(M)** Cell volume determined using Fluorender on wholemount images of EDL (n=3-7 mice/DPI; n= 14-40 SCs/DPI). **(N)** Length of all cellular projections, **(O)** average individual projection length, **(P)** distribution of SCs with 1-4 projections, and **(Q)** distribution of SCs with 1-6+ terminal ends of GFP+ SCs in uninjured or Galert state (n=7 mice/DPI; n=36-42 cells/DPI). **(R)** Experimental design for labeling proliferative SCs via EdU 4h prior to harvest. **(S-T)** Representative wholemount images of EdU+ GFP+ SCs at 1, 1,5, and 2 DPI **(S)**, quantification of EdU+GFP+ SCs **(T)**, and representative GFP+ dividing SCs **(U)** (n=3-4 mice/DPI; n=547-2664 GFP+ SCs/DPI). **(V-Y)** Division angles of dividing Edu+ GFP+ SCs 1-2 DPI. **(V)** Model of possible SC division angles; cell (a) represents planar divisions, cell (b) apical-basal, and cell (c) typical cell observed in vivo. **(W-Y)** Quantification of SC division angles. Red dotted line indicates minimum X-Z division angle for designation of apical-basal division (n = 3-4 mice/DPI and 8-49 EDU+GFP+ SCs measured/DPI). Scale bars = 50um. Histogram values are mean ± SEM; * p < 0.05, ** p < 0.01, *** p < 0.001 and **** p < 0.0001.

BaCl_2_ injury led to a transient increase, subsequent decline, and then recovery in muscle mass by 21 DPI (Fig.1B). To assess the overall effect of injury on SCs and their progeny we analyzed the number of mononuclear myogenic cells. We found by FACS and quantification of GFP+ myogenic cells in wholemount images (via the image rendering software Fluorender) that there was a rapid increase in mononuclear myogenic cells between 2.5 and 3 DPI (Fig. 1C-E). High numbers of mononuclear myogenic cells were present through 14 DPI and then declined as myogenic cells fused to make myofibers or returned to quiescence (Fig. 1C-D). Given this overall framework for muscle recovery after acute damage, we then proceeded to analyze in whole mount the cellular processes by which SCs regenerate myofibers.

### Quiescent, Galert, and newly activated SCs are distinct in morphology and size

During homeostasis, quiescent SCs occupy a distinct niche between the myofiber sarcolemma and the BM ^5, 12, 44, 45^. To visualize quiescent SCs in vivo, we imaged perfused fixed wholemount uninjured EDLs by confocal microscopy and reconstructed their three-dimensional structure with Fluorender (Fig. 1F-K, Video S1) ^46^. In these muscles, TOM labeled all uninjured myofibers and highlighted the regular banding of sarcomeres, present at the expected 2.5um interval (Fig. 1G-H) ^47^. Capillaries lying between myofibers were also TOM+. Other cell types, such as fibroadipogenic progenitors (FAPs), were labeled at much lower levels with TOM from the *Rosa^mTmG^* reporter than myofibers and capillaries and therefore are not shown. We found that GFP+ quiescent SCs resided between the sarcolemma and the BM (shown via laminin immunofluorescence), often in close proximity to capillaries (arrows in Fig. 1G-I, Video S1; see also ^41, 48^). SCs were elongated, generally parallel to the long axis of the myofiber and spanned 20-25 sarcomeres (Fig. 1G-H). Strikingly, individual SCs possessed long cellular projections that branched into numerous terminal ends (Fig. 1N-Q). Although quiescent satellite cells have generally been depicted as fusiform cells with little cytoplasm ^14, 49, 50^, an electron microscopy study ^51^ and recent in vivo studies ^41, 52–55^ describe such projections. Our findings demonstrate that our imaging and visualization techniques are sensitive enough to preserve these delicate structures and also provide further evidence that in vivo quiescent SCs have a distinctive morphology, with multiple long projections.

SCs have been identified that are neither fully quiescent nor activated, but rather primed for more rapid activation, in a state termed Galert ^56^. We examined Galert SCs (as defined by ^56^) to test whether cellular morphology was changed in this state. Previous studies ^56^ reported that Galert SCs are larger than quiescent SCs. However, we found that by FACS Galert SCs were significantly smaller in diameter than quiescent SCs (Fig. 1L), although no difference in volume was detected when measured via Fluorender analysis of confocal images (Fig. 1M). We observed no difference in the number of projections or terminal ends in Galert SCs (Fig. 1P-Q). However, the length of the cellular projections was shorter (Fig. 1N-O; and similar to findings of ^54, 55^) and is consistent with their being primed for activation.

Upon activation, SCs displayed a greatly altered morphology by 1 DPI. SCs were significantly enlarged with maximal cell size reached by 1.5 DPI (Fig. 1L-M). The distinctive thin projections of quiescent SCs were absent and instead replaced with broader lamellipodia (Fig. 1F-H), indicative of motile cells ^52, 57^. Notably, all activated SCs were found confined to the inside of residual BM tubes (termed "ghost fibers" by ^52^) although the TOM+ sarcolemma and sarcomeres of the injured myofibers were now degraded and largely absent (Fig. 1F-K). Activated SCs continued to generally be in close proximity to capillaries (arrows, Fig 1G-I).

### SCs proliferate via planar divisions

After activation, SCs proliferate to provide myoblasts and myocytes necessary to regenerate myofibers and also to self-renew quiescent SCs. As noted by others (e.g. ^56, 58^), the initial round of cell division in vivo occurs approximately 30 hours after acute injury, while all subsequent divisions occur in roughly half that time. Consistent with this, the number of GFP+ myogenic cells expanded between 2 and 3 DPI (Fig. 1C-E). Of considerable interest is whether the early cell divisions are asymmetric (i.e. a SC gives rise to one SC and one committed myoblast) or symmetric (i.e. a SC gives rise to two SCs) ^12^. The niche within which quiescent SCs reside is inherently asymmetric, with the BM establishing the basal surface and the sarcolemma providing the apical surface ^5, 12, 44^. Several studies ^11, 59, 60^, based on cultured myofibers with attached SCs, have suggested that early asymmetric divisions are manifested as apical-basal cell division events, with the daughter cell adjacent to the BM becoming a quiescent SC, while the daughter adjacent to the sarcolemma differentiates into a MyoD+ myoblast. However, it is unclear whether these in vitro observations are applicable to the situation in vivo. To test for the frequency of apical-basal divisions in vivo, we injected mice with a pulse of EdU 4 hours prior to harvest (Fig. 1R) and analyzed the division angles of EdU+ GFP+ myogenic cells captured in the process of division (identified as pairs of SCs which share an intensely GFP+ membrane, are condensed in shape, and with nuclei closer than 20 um) at 1, 1.5 and 2 DPI (arrows, Fig. 1S). At all timepoints, we found no apical-basal divisions, but rather planar cell divisions with the two daughter cells aligned parallel to the long axis of the damaged myofiber (Fig. 1U-Y). Thus, we find no support in vivo for apical-basal divisions as a widespread mechanism for generating asymmetric cell fates.

### Myogenic cells proliferate and accumulate randomly within narrowing residual BM tubes 2-3 DPI

Myogenic cells continue to undergo dramatic morphological changes 2-3 DPI. At 2 DPI GFP+ myogenic cells become flattened (Fig. 2A-B, Video S2) and reside adjacent to the inner side of the BM (Fig. 2D-E). This flattened morphology appeared to be dictated by the presence of residual sarcomeric proteins which filled the interior space of BM tubes (Fig. 2H). By 2.5-3 DPI, as sarcomeric debris was removed, the myogenic cells were no longer restricted to the periphery of the tubes and became less flattened and more compact in shape (Fig. 2A-E).

**Figure 2.**
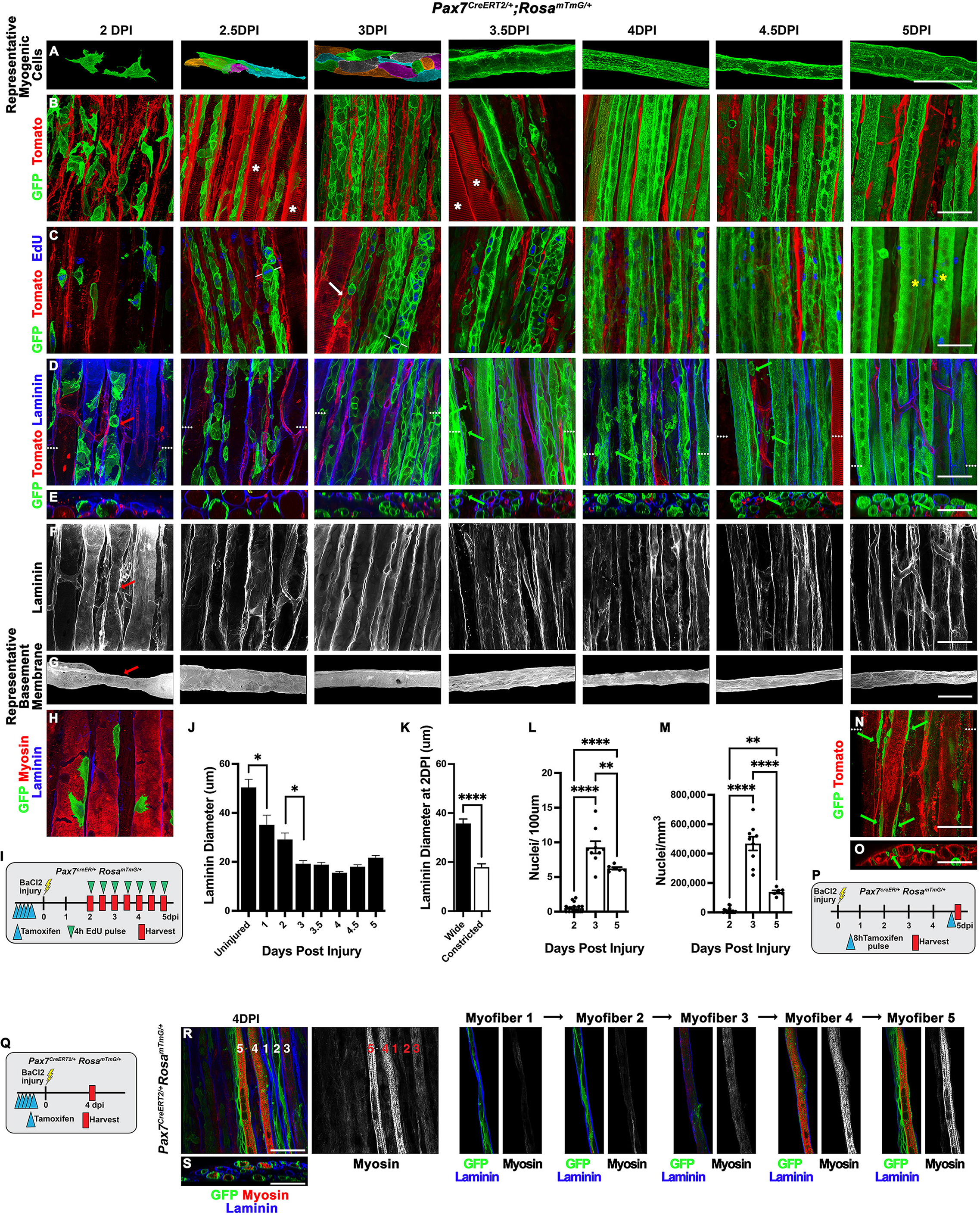
Regenerated myofibers form via a wave of density-dependent fusion 3.5-4.5 DPI followed by alignment of myonuclei by 5 DPI. **(A-G)** Representative wholemount images of EDL muscles 2-5 DPI (n= ≥ 3 mice/DPI). **(A)** Representative segmented GFP+ myogenic cells or nascent myofibers. Individual cells pseudo-colored at 2.5-3.0 DPI. **(B)** Superficial view of GFP+ myogenic cells and TOM+ uninjured myofibers with intact sarcomeres (*’s at 2.5 and 3.5 DPI), and TOM+ capillaries (all columns). **(C)** EdU labeling (in blue) 4h prior to harvest. At 2.5 DPI white lines show division angle of dividing myogenic cell. At 3 DPI white arrow highlights junction of damaged TOM+ myofiber being repaired by GFP+ myogenic cells. At 5 DPI only peripheral myonuclei are labeled (yellow *’s) **(D)** Laminin outlines residual BM tubes and surrounds capillaries. Dotted white lines indicate level of cross-sections shown in **(E)**. Green arrows in (D-E) mark interstitial GFP myogenic cells residing outside BM tubes. **(F)** Superficial view with only laminin of scans in (D). **(G)** Representative laminin+ BM tubes. Red arrows in (D, F, G) show a necked BM tube. **(H)** At 2 DPI GFP+ myogenic cells are confined to BM tubes in regions where degrading, non-sarcomeric myosin (in red) is absent. **(I)** Experimental design for genetic lineage tracing of SCs and EdU labeling, 2.5-5.0 DPI. **(J)** Diameter of laminin+ BM tubes of uninjured and injured myofibers 1-5 DPI (n=2-3 mice/DPI; n=4-6 laminin+ tubes/DPI; n=5 measurements/tube). **(K)** Comparison of wide and narrow diameters of necked laminin+ BM tubes at 2 DPI (n=3 mice; n=11 tubes). **(L-M)** Number of myonuclei/100um length (L) or volume (M) of regenerating myofiber (n=3 mice/DPI; 3-8 myofibers/DPI). (**N-P**) Single TAM pulse 8h prior to 5 DPI harvest (scheme in P) shows GFP+ myogenic cells (green arrows, N-O) contribute and fuse peripherally to nascent myofibers. Dotted white lines in (N) indicate level of cross-section in **(O)**. **(Q-S)** Nascent GFP+ myofibers labeled (experimental scheme, Q) immunolabeled with perinatal myosin show that formation of centralized chains is coincident with development of organized sarcomeric myosin; representative progression from newly formed myofiber (1) to most mature myofiber (5). All images scale bars = 50um. Histogram values are mean ± SEM; * p < 0.05, ** p < 0.01, *** p < 0.001 and **** p < 0.0001.

During 2-3 DPI myogenic cells are highly proliferative (Fig. 1T, 2C). Previous reports ^34, 61^ hypothesize that this proliferative phase is critical to the generation of the chains of aligned, centralized myonuclei characteristic of regenerated myofibers. In particular, it was proposed that these chains result from aligned successive planar divisions of myogenic cells. Although our quantification of division angles revealed that most cell divisions are planar (Fig. 1U-Z, white lines at 2.5 and 3 DPI in Fig. 2C), to our surprise we found that GFP+ myogenic cells were randomly distributed within the BM tubes at this stage (Fig. 2B-D, Video S2). This disproves the hypothesis that centralized chains are a simple consequence of successive planar divisions and indicates that such chains must arise from another mechanism.

These images also provide other insights. First, throughout the first 3 days of regeneration, we found that all myogenic cells were sequestered within the BM tubes. This indicates that SCs do not travel outside their associated BM during this early phase, and therefore initial regeneration of individual damaged myofibers is mediated by the SCs associated with them. Second, with a robust BaCl_2_ injury most regenerating myofibers were composed entirely of GFP+ myogenic cells (indicative of myofibers regenerating de novo from SCs), but we also observed hybrid myofibers composed of original TOM+ myofiber stumps flanked by EdU+ GFP+ proliferating myoblasts (white arrow, Fig. 2C, Video S2). Thus integration of old damaged and de novo myofibers occurs.

During this time the residual BM tubes underwent significant morphological changes (Fig. 2D-G, Video S2). In uninjured myofibers, the laminin+ tubes averaged 50 um in diameter (Fig. 1J-K, 2J) but shrank to 20 μm by 3 DPI (Fig. 1J-K, 2F-G, J). Particularly at 2 DPI, “necked” tubes with wide and constricted regions were visible (red arrows Fig. 2D, F,-G, quantified in 2K, Video S2); these are snapshots of BM tubes caught in the contraction process. We hypothesize that this contraction results from the clearance of sarcomeric debris, such that the constricted regions are cleared of debris while the wide regions still retain debris (Fig. 2H).

### Density-dependent myocyte-myocyte fusion re-establishes myofibers 3.5-4.5 DPI with chains of centralized nuclei subsequently appearing as sarcomeres form

Between 3.5 and 4.5 DPI regeneration takes a dramatic turn, as a wave of fusion resulted in the re-establishment of most myofibers by 5 DPI (Fig. 2 A-E, Video S2). The number of GFP+ myogenic cells increased rapidly so that their number was maximal at 3-4 DPI (Fig. 1C-D, 2C). Concurrent with this increase in cells, the BM tubes shrank to their smallest diameter at 3-4.5 DPI (Fig. 2J). As a consequence of this rapid increase in myogenic cells and decrease in volume of the tubes in which they reside, myogenic cells reached their maximal density at 3-4 DPI (Fig. 2L-M). An increase of myogenic cell density to 500,000 cells/mm^3^ (Fig. 2L-M) appears to trigger a massive wave of density-dependent fusion. Because fusion appeared to occur nearly synchronously along the length of the regenerating myofibers, myocyte-myocyte fusions likely dominate this process rather than myocyte to myofiber fusion.

By 5 DPI all regenerated myofibers contain centralized chains of regularly-spaced compact nuclei. However, this is not a feature of newly regenerating myofibers at 4-4.5 DPI (Fig. 2A-D, Video S2). Analysis of tile scans (Fig. S2A-B) of whole EDLs at 4.5 DPI showed the ubiquitous presence of newly-fused, thin multinucleate myofibers, but chains of regularly-spaced compact myonuclei were present only in some regions of these myofibers (asterisks in Fig. S2B). This indicates that nuclear chains form secondarily to fusion. We first hypothesized that the centralized chains may form in response to reinnervation of the myofibers. To test this, we transected and retracted the left common peroneal nerve to prevent reinnervation and also injured the left EDL with BaCl_2_ (Fig. S2C). Denervation was successful as the EDL mass decreased significantly (Fig. S2D). However, centralized chains of regularly spaced nuclei were still apparent at 7 DPI, demonstrating that innervation is not required for their formation. Instead, we found that chains of regularly-spaced nuclei arose coincident with the periodic banding in GFP+ myofiber membranes that is indicative of sarcomere formation (Fig. 2D, 5 DPI), suggesting that the two processes are linked. Indeed, detailed comparison of myofibril and sarcomere formation (via analysis of myosin heavy chain immunofluorescence) and myonuclei showed that chains of regularly spaced nuclei only appeared in newly regenerated myofibers that have myofibrils with sarcomeric banding (Fig. 2Q-S, Video S2). These observations suggest that the myonuclei are corralled into centralized chains of regularly-spaced compact nuclei by the formation of the myofibrils and sarcomeres in which they are embedded.

### Myofibers enlarge by myocyte-myofiber fusion and addition of peripheral myonuclei

Myofibers are regenerated by 5DPI, but regenerated myofibers are about half the diameter of an uninjured myofiber (Fig. 3D). Between 5 and 14 DPI, myofibers enlarged (Fig. 3A-D, Video S3) and the number of myonuclei increased through at least 21 DPI (Fig. 3E). This suggests that SCs continued to contribute to nascent myofibers for many days after the initial wave of fusion. To test this, we gave *Pax7^CreERT^*^2^*^/+^;Rosa^mTmG/+^* mice a single dose of TAM at various days post injury, harvested muscles at 14 DPI, and quantified numbers of GFP+ and TOM+ regenerated myofibers with centralized nuclei (Fig. 3F-I, M). As a control, we gave some mice a single dose of TAM and harvested uninjured EDLs at 14 DPI. As expected, a few uninjured myofibers were GFP+, indicating that SCs contributed to these homeostatic myofibers (Fig. 3M; see also ^62^). As another control, we gave mice a single dose of TAM prior to injury and harvested EDLs at 14 DPI (Fig. 3G). This resulted in 84% of regenerated myofibers being GFP+ (Fig. 3M) as compared with nearly 100% GFP+ regenerated myofibers when 5 TAM doses were given prior to injury. When one dose of TAM was given at 1 DPI, nearly all regenerated myofibers were GFP+ (Fig. 3G, M). Single TAM doses given at later days post-injury resulted in progressively fewer GFP+ regenerated myofibers (Fig. 3H, I, M). TOM+ regenerated myofibers in these experiments were derived from earlier SCs that had not been labeled with GFP when TAM was delivered. Surprisingly, few regenerated myofibers were GFP+ when TAM was delivered at 7 DPI. We reasoned that the efficiency of the *Pax7^CreERT^*^2^*^/+^* allele was lower with only a single dose of TAM and so repeated the single-dose TAM experiments at 7 and 10 DPI experiments with *Pax7^CreERT^*^2^*^/CreERT^*^2^*;Rosa^mTmG/+^* mice (Fig. 3J-L, N). With two copies of the Cre, 50% of regenerated myofibers were GFP+ when TAM was delivered at 7 DPI. Even with two copies of Cre, few regenerated myofibers were GFP+ when TAM was delivered at 10 DPI, indicating that few SCs detectably contributed to regeneration of myofibers at this timepoint, even though the number of myonuclei slowly increased until at least 21 DPI (Fig. 3E). Overall, these data indicate that the major phase of SC contribution to regeneration of myofibers is complete by 10 DPI.

**Figure 3.**
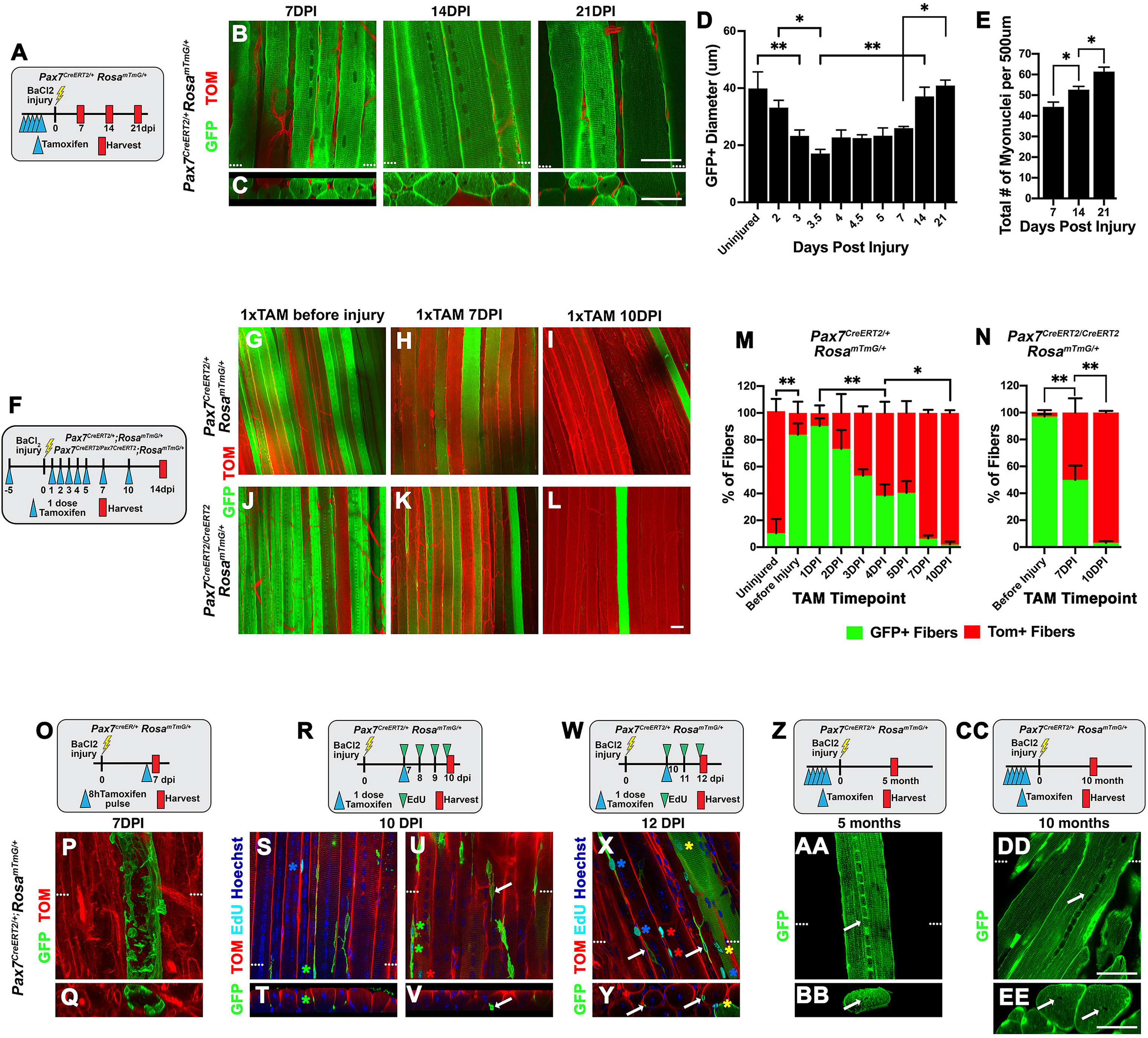
Myofibers enlarge by contribution of myogenic cells to peripheral nuclei, with centralized chains generated by 5 DPI persisting for months after regeneration. **(A)** Experimental scheme for labeling SC-regenerated myofibers 7-21 DPI. **(B-C)** Representative wholemount images of EDL muscles 7, 14, and 21 DPI (n= ≥ 3 mice/DPI). Dotted lines in (B) show level of cross-section in (C). **(D)** Quantification of diameter of uninjured and GFP+ regenerated myofibers (n=3-6 mice/DPI; n= 73-304 GFP+ myofibers/DPI; n= 3measurements/myofiber). **(E)** Quantification of myonuclei/500 um length (n= 3 mice/DPI; n = 18 myofibers/DPI). Experimental scheme **(F)**, representative wholemount images of EDLs from *Pax7^CreERT^*^2^*^/+^;Rosa^mTmG/+^* or *Pax7^CreERT^*^2^*^/CreERT^*^2^*;Rosa^mTmG/+^* mice **(G-L)**, and quantification **(M-N)** of SC contribution when labeled at different DPI and assayed at 14 DPI (n=2-5 mice/DPI; n=50-214 myofibers/DPI). **(O-Q)** GFP+ SCs contribute to peripheral region of nascent myofibers at 7 DPI. **(R-Y)** SCs at 10 DPI contributing (green asterisks, S-U) or at 12 DPI contributed (yellow asterisks, X-Y) peripheral EdU+ GFP+ myonuclei. A few EdU+ myonuclei are GFP-(red asterisk, U, X) and EdU+ non-myogenic nuclei (blue asterisk, S, X) are also present. SCs with quiescent morphology are present (white arrows, U-V, X-Y). **(Z-EE)** Chains of centralized myonuclei (white arrows) persist in regenerated myofibers 5 and 10 months post injury. Dotted white lines in (P, S, U, Y, X, AA, DD) show level of cross-section in (Q, T, V, Y, CBB, EE). Images A-EE, scale bars = 50um.

Given that centralized chains of myonuclei are present by 5 DPI, we next tested the location of myonuclei contributed by SCs at 5 and 7 DPI. To test this, we gave *Pax7^CreERT^*^2^*^/+^;Rosa^mTmG/+^* mice a single dose of TAM 8h prior to harvesting EDLs at 5 or 7 DPI (Fig. 2N-P, 3O-R). The majority of regenerated myofibers were TOM+, as they were generated by SCs prior to TAM. Elongate GFP+ myogenic cells (but without the thin projections distinctive of quiescent SCs) were found fusing peripherally to regenerating TOM+ myofibers at 5 DPI (green arrows, Fig. 2N-P, Video S2). At 7 DPI we captured GFP+ myogenic cells fusing to form a peripheral scaffold around a regenerated myofiber (Fig 3P-Q). These data indicate that SCs fusing to regenerating myofibers after 4 DPI contribute peripheral myonuclei. In addition, labeling with EdU 4 h prior to harvest demonstrated that at 5 DPI all EdU+ myonuclei were peripheral (yellow asterisks, Fig. 2C, Video S2) and no myonuclei in the centralized chains were EdU+. Furthermore, EdU administration later than 5 DPI (at 10 or 12 DPI; Fig. 3R-Y) never labeled myonuclei in the centralized chains, but labeled only recently adding (green asterisks marking EdU+GFP+ elongate myogenic cells, Fig. 3S-U, Video S3) or added peripheral myonuclei (yellow asterisks marking EdU+ myonuclei in GFP+ myofibers, Fig. 3X-Y). Note that some EdU+ myonuclei were TOM+, indicating that these myonuclei were derived from recently dividing myogenic cells that were Pax7-2 or 3 days before harvest and so unlabeled by TAM (red asterisks, Fig. 3U, X). Overall, these data indicate that after formation of centralized chains by 4.5 DPI all subsequently added myonuclei are located peripherally. Interestingly, these later contributing SCs may not only participate in regeneration of the damaged myofibers with which they were associated, but may also contribute to regeneration of other – presumably nearby – myofibers, as SCs residing in the interstitial space between BM tubes were first observed at 3.5 DPI and common at subsequent timepoints (green arrows, Fig. 2D, Video S2) .

### SCs with quiescent morphology appear by 10 DPI

There has been considerable debate about when SCs return to quiescence and repopulate their niche between the myofiber’s BM and sarcolemma ^11, 59, 60, 63–65^. We have found that quiescent SCs are morphologically unique, typically with 2 thin projections (Fig. 1F-H, N-Q). Our experiments labeling all SCs and their derivatives via TAM prior to injury did not allow us to readily identify the re-appearance of quiescent SCs because of the high density of GFP+ myogenic cells throughout the regenerating muscle. Instead, we searched for the first presence of morphologically distinct quiescent SCs by a single dose of TAM prior to harvest (either 8 h prior to harvest at 5 or 7 DPI or 48 h prior to harvest at 10 or 12 DPI). We readily found morphologically quiescent SCs that were EdU-(see EdU labeling strategies in Fig. 3 R, W) at 10 and 12 DPI (white arrows, Fig. 3U-Y, Video S3). However, we did not identify morphologically distinctive quiescent SCs at 5 or 7 dpi. Although we cannot exclude that molecularly distinct quiescent SCs are present earlier, our data indicate that morphologically distinct quiescent SCs do not appear until after 7 DPI.

### Centralized chains of myonuclei persist in regenerated myofibers for months

While centralized chains of myonuclei have been recognized as a hallmark of regenerating myofibers for more than 75 years ^32–35^, it has been unclear how persistent these chains are in regenerated myofibers. In development ^66^ and in culture ^67^, centralized myonuclei have been found to transition to peripheral nuclei. Therefore, it has been assumed by many ^36, 37^ that the chains of centralized myonuclei eventually migrate peripherally. However, a more recent study ^33^ indicates that centralized nuclei persist for nearly two years, although this study was limited by the lack of a method to establish which individual myofibers were regenerated independent of the presence of centralized nuclei. Our genetic labeling of SCs and their derivatives now allow us to tag regenerated myofibers and determine the fate of the centralized chains of myonuclei. We gave TAM to *Pax7^CreERT^*^2^*^/+^;Rosa^mTmG/+^* mice, injured EDLs, and harvested EDLs 5 or 10 months post injury (Fig. 3Z, CC). As expected, most myofibers were GFP+, indicating that they were regenerated. Strikingly, we found that at 5 and 10 months post-injury nearly every GFP+ regenerated myofiber contained regions with centralized chains of myonuclei (Fig. 3AA-BB, DD-EE). Quantification of a subset of samples stained with Hoechst revealed that at 10 months 98% of GFP+ regenerated myofibers had chains of centralized nuclei (n = 47/48 myofibers in 2 mice). This indicates that, unlike development in vivo or myogenesis in vitro, centralized nuclei that form as the result of regeneration do not migrate to the periphery and therefore indelibly mark regenerated myofibers.

### Regeneration differs from development of myofibers

Muscle development and regeneration are often suggested to use similar molecular and cellular processes ^39, 40^. However, development and regeneration have different temporal and functional constraints. In mice, limb myogenesis occurs in three extended phases: embryonic myogenesis establishes the basic muscle pattern (E10.5-14.5), fetal myogenesis enlarges myofibers and begins to establish distinct contractile fiber types (E14.5 – E18.5), and several weeks of postnatal myogenesis are important for muscle growth and maturation ^39, 68, 69^. The first two phases occur within the protective confines of the pregnant mother and so there are no functional demands on developing muscle. In contrast, we find that after BaCl_2_ destruction of myofibers, myofibers are regenerated by 4.5 DPI. Presumably this accelerated formation of myofibers during regeneration is necessitated by the functional requirements of an ambulatory mouse. Altogether these observations indicate that myogenesis during development and regeneration may be similar overall, but key differences are likely. We therefore compared our whole mount in vivo findings on regeneration with developmental myogenesis.

We examined the TA at E12.5, when embryonic myogenic cells fuse to make primary myofibers, and at E14.5, when fetal myogenic cells both fuse to primary myofibers and fuse to one another to make secondary myofibers ^68, 69^. We first examined whether the BM was essential for myofiber development. We examined E12.5 *Pax3^Cre/+^;Rosa^mTmG/+^* embryos, in which *Pax3^Cre^* genetically labels all embryonic muscle progenitors and their myogenic derivatives in the limb ^69^. At this stage myogenic cells were fusing to make myofibers with continuous GFP+ membranes (white arrows, Fig. 4A, Video S4). However, in contrast to regeneration after BaCl_2_ injury, this fusion happened in the absence of a continuous laminin+ BM. Instead, the BM was patchy, appearing as small dots adjacent to the newly formed myofiber sarcolemma (white arrows, Fig. 4A-B, Video S4). We also examined E14.5 *Pax7^Cre/+^;Rosa^mTmG/+^* embryos, in which *Pax7^Cre^* labels all fetal muscle progenitors and their myogenic derivatives in the limb ^69^. The myofibers of the TA were larger in diameter (white arrows, Fig. 4C, Video S4) than at E12.5 and parts of the myofibers have areas of continuous laminin+ BM (white arrows, Fig. 4C-D, Video S4). Thus, unlike regeneration, a continuous BM is absent during primary myogenesis, but as development proceeds a BM builds up to surround all myofibers and is present during fetal myogenesis.

**Figure 4.**
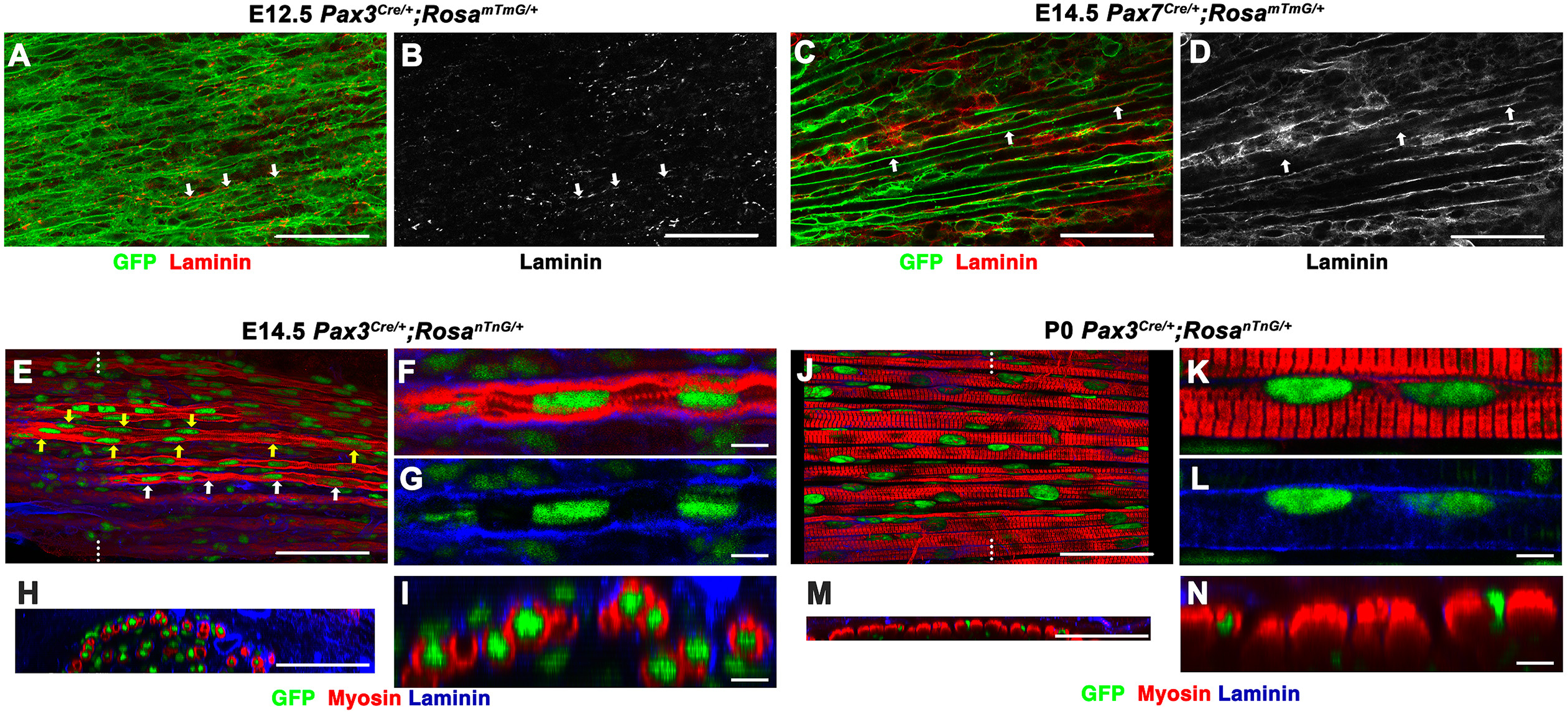
Development of myofibers is distinct from regeneration; myogenesis begins in the absence of continuous BMs and initially centralized myonuclei move peripherally. **(A-B)** Representative images of TA at E12.5 of *Pax3^Cre/+^;Rosa^mTmG/+^* mice showing GFP+ forming myofibers with speckly laminin (white arrows). **(C-D)** Representative images of TA at E14.5 of *Pax7^Cre/+^;Rosa^mTmG/+^* mice showing areas of continuous laminin adjacent to GFP+ sarcolemma (white arrows show one myofiber with no laminin on left side, but laminin on right side). **(E-J)** Representative images of *Pax3^Cre/+^;Rosa^nTnG/+^* mice at E14.5 (E-I) or P0 (J-N) showing primary myofibers with single row of centralized nuclei (white arrows, E) or secondary myofibers with several rows of nuclei (yellow arrows, E). Myonuclei are peripheral by P0 (J-N). (H) and (M) are cross-sections of (E) and (J), respectively; location of cross-sections shown as white lines on (E) and (J). (F-G), (I), (K-L), and (N) are magnifications of (E), (H), (J), and (M), respectively. Images A-E, H, J, and M scale bars = 50um; Images F-G, I, K-L, and N scale bars = 5um.

We also examined the location of myonuclei in the developing myofibers, as it has been proposed that nascent myofibers have centralized nuclei that move peripherally as the myofibers mature ^37, 38^, although this model is largely based on in vitro studies. We examined at E14.5 *Pax3^Cre/+^;Rosa^nTnG/+^* embryos, in which all myogenic nuclei are GFP+, that were also labeled with antibodies to laminin and perinatal myosin (Fig. 4E-I, Video S4). At this stage there were primary myofibers with a single row of dispersed myonuclei located centrally and surrounded on most sides by irregular, partially formed myofibrils (white arrows, Fig. 4E and 4F-I). There were also some secondary myofibers (yellow arrows Fig. 4E) which contained more than one row of myonuclei, and the additional nuclei were peripherally located (left yellow arrows Fig. 4E) and likely the result of recently fused myocytes. It is notable that the organization of the E14.5 myonuclei is distinct from the centralized myonuclei of regenerated myofibers, which form centralized chains of regularly spaced, compact myonuclei. We also examined the TA at P0 and found that the myofibers were considerably larger and the myofibrils were aligned in distinct sarcomeres (Fig. 4J-N, Video S4). Notably, all myonuclei of these myofibers were located peripherally, outside the myofibrils and beneath the BM. Therefore, unlike regeneration, as developing myofibers grow in diameter, their myonuclei shift to a peripheral position.

### Macrophages are required for proper density-dependent fusion of myogenic cells

Fusion of myogenic cells at 3.5-4.5 DPI is essential for rapidly re-establishing myofibers, and our data suggest that this event is density-dependent. This increase in myogenic cell density is a consequence of increasing numbers of myogenic cells that are contained within a shrinking BM, which reaches a minimal size during this critical time. We hypothesized that this BM shrinkage is due to the removal of myofibril debris within the BM tubes by macrophages. To test this, we experimentally depleted macrophages during regeneration.

During regeneration, macrophages have many critical functions, including orchestration of the inflammatory response, clearance of cellular debris (efferocytosis), regulation of SC proliferation and fusion, and regulation of FAPs ^70–73^. We found that macrophages were recruited to muscle, reached peak numbers at 3 DPI (Fig. 5A), and migrated inside the BM tubes in close contact with myogenic cells (white arrows, Fig. 5B, Video S5). To test whether macrophage clearance of debris is required for BM shrinkage, we depleted macrophages at different time points via intraperitoneal delivery of clodrosome, a liposome-encapsulated clodronate, which kills phagocytic macrophages (Fig. 5C; ^74^). Delivery of encapsome, liposomes lacking clodronate, served as a negative control.

**Figure 5.**
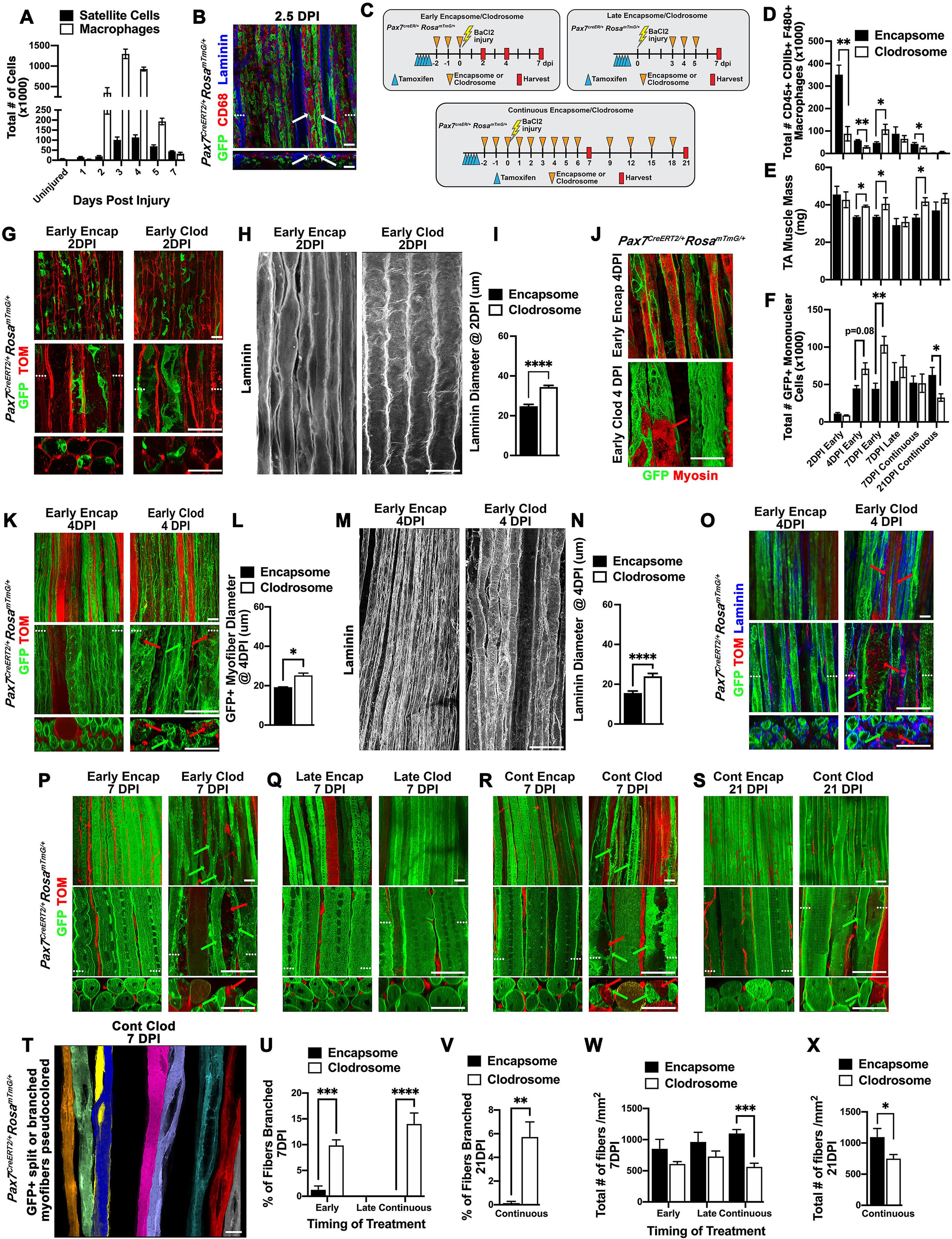
Macrophages are essential to clear debris, enabling density-dependent myogenic cell fusion and preventing formation of branched myofibers. **(A)** Quantification by FACs of macrophages and SCs during regeneration (n = 2-4 mice/DPI). **(B)** At 2.5 DPI macrophages (labeled by CD68) have penetrated the BM tubes and reside adjacent to GFP+ myogenic cells (white arrows). **(C)** Experimental schemes for macrophage depletion. **(D-F)** At different DPIs and with early, late, or continuous clodrosome (or control encapsome) administration to TAs, quantification via FACS analysis of macrophages (D), TA mass (E), or via FACS GFP+ mononuclear myogenic cells (F) (n= 2-7 mice/condition). **(G-I)** Macrophage depletion does not affect SC activation, but prevents normal BM shrinking. (n= 3 mice/condition; n=51-53 myofibers/condition) **(J-O)** Macrophage depletion leads to persistence of disorganized myosin debris (red arrow, J) and TOM+ sarcolemma debris (red arrows, K and O), poorly fused (green arrows, K and O) and wider GFP+ myogenic nascent myofibers (L) (n=3 mice/condition; 65-83 myofibers/condition), and persistently wide BM tubes (M, N) (n= 3 mice/condition; n=25-27 myofibers/condition) at 4 DPI. **(P-S)** Early (P) or continuous (R-S) macrophage depletion causes aberrantly fused and branched myofibers (green arrows) with persistent TOM+ sarcolemma debris (red arrows), but such an effect is not seen with late clodrosome administration. **(T-V)** Macrophage depletion leads to many branched myofibers (pseudo-colored in T and quantified in U-V) (n=3-6 mice/condition; 197-655 myofibers/condition). **(W-X)** Continuous clodrosome administration leads to regeneration of fewer myofibers (n=5-6 mice/condition; n=387-646 myofibers/condition). For all encapsome or clodrosome experiments, n = ≥ 2 mice/experiment. In all whole mount images with three panels dotted white lines in middle panel show level of cross-section in lower panel. Scale bars = 50um. Histogram values are mean ± SEM; * p < 0.05, ** p < 0.01 *** p < 0.001, and *** p < 0.0001.

Delivery of clodrosome prior to injury (Fig. 5C) resulted in a substantial depletion of macrophages in the muscle by 2 DPI (Fig. 5D). Macrophage depletion did not result in any change in muscle mass, number of GFP+ mononuclear cells, or cellular signs of SC activation at 2 DPI (Fig. 5D-G). However, the diameter of BM tubes remained wider with macrophage depletion (Fig. 5H-I). At 4 DPI, muscles in which clodrosome was delivered prior to injury had a significant reduction in macrophages (Fig. 5D) and muscles were heavier (Fig. 5E). In control muscles, most myofibers were re-established by the wave of density-dependent fusion (Fig. 5K, O, Video S5). In contrast, clodrosome treatment resulted in wider GFP+ myofibers with many unfused or incompletely fused myogenic cells (green arrows, Fig. 5K, O; quantified in Fig. 5L) and wider BM tubes (Fig. 5M-N). Within these wider myofibers were fragments of residual, un-phagocytized TOM+ sarcolemma from the damaged myofibers (red arrows, Fig. 5K, O). Also, while newly formed sarcomeres were present in nascent myofibers in control muscles (Fig 5J), myofibers with residual disorganized myosin debris were frequently seen in clodrosome-treated muscles (red arrow, Fig. 5J). This physical impedance of un-phagocytosed sarcomeric proteins and cell membranes led to poor or aberrant fusion of myogenic cells, resulting in regenerated myofibers with wide gaps (green arrows, Fig. 5K, O, Video S5). Consistent with this, the number of mononuclear, unfused GFP+ myogenic cells was somewhat elevated (p = 0.08, Fig. 5F).

Overall, these experiments demonstrate that phagocytosis by macrophages is critical for clearance of necrotic debris, allowing BM tubes to contract and density-dependent fusion to proceed unimpeded.

### Depletion of M1 macrophages during regeneration leads to split and branched myofibers

To see whether macrophage depletion had later effects on muscle regeneration we analyzed muscles at 7 DPI. Macrophage depletion via clodrosome treatment prior to injury caused 10% of regenerated myofibers to have an aberrant morphology; myofibers were split or branched (green arrows, Fig. 5P; pseudo-colored, Fig. 5T; quantified, Fig. 5U) and some branched myofibers still retained uncleared TOM+ debris (red arrows, Fig. 5P). We also tested whether continuous delivery of clodrosome would exacerbate the phenotype (Fig. 5C). While we found that with continuous clodrosome delivery 14% of regenerated myofibers were branched (green arrows, Fig. 5R; Video S5; quantified, Fig. 5U) often surrounding residual debris (red arrows, Fig. 5R), there was no statistical difference in the number of branched myofibers between early and continuous clodrosome delivery (Fig. 5U). In addition, continuous clodrosome delivery led to the regeneration of fewer myofibers (Fig. 5W). We also attempted to deplete macrophages at a later time during regeneration (daily delivery at 3-5 DPI; Fig. 5C), but at 7 DPI macrophages were not significantly depleted and the morphology of myofibers was similar to control myofibers (Fig. 5D, Q). Nevertheless, comparison of early and continuous clodrosome treatments argues that depletion of early macrophages was the dominant cause of the phenotype. Early treatment of muscle with clodrosome (2 and 1 days before injury and day of injury) led to depletion of macrophages at 2 and 4 DPI, but not at 7 DPI (macrophages were actually upregulated compared to controls, suggesting some compensation for previous depletion), while continuous treatment of muscle led to depletion of macrophages through 7 DPI (Fig. 5D). The similarity in the branching phenotype with early and continuous depletion of macrophages strongly suggests that it is depletion of the early M1 pro-inflammatory macrophages that is the cause of this phenotype.

We also tested whether depletion of macrophages led to a persistent phenotype at 21 DPI. With continuous clodrosome delivery, macrophages were effectively depleted at 7 DPI, although no discernable difference was detected at 21 DPI (when macrophages return to low basal levels) (Fig. 5D). Nevertheless, we found that macrophage depletion led to persistent elevated levels of branched myofibers (Fig. 5T, V) and also decreased numbers of regenerated myofibers (Fig. 5X).

## DISCUSSION

SCs are well established to be the stem cells necessary and sufficient for regeneration of myofibers ^6–8^, but how regeneration of the three-dimensional architecture of linear and aligned myofibers occurs is largely unknown. By genetically labeling SCs, imaging, and reconstructing regeneration in three dimensions, our study has provided surprising new insights and leads to a new synthetic model (Fig. 6) of how SCs quickly re-establish muscle structure.

**Figure 6.**
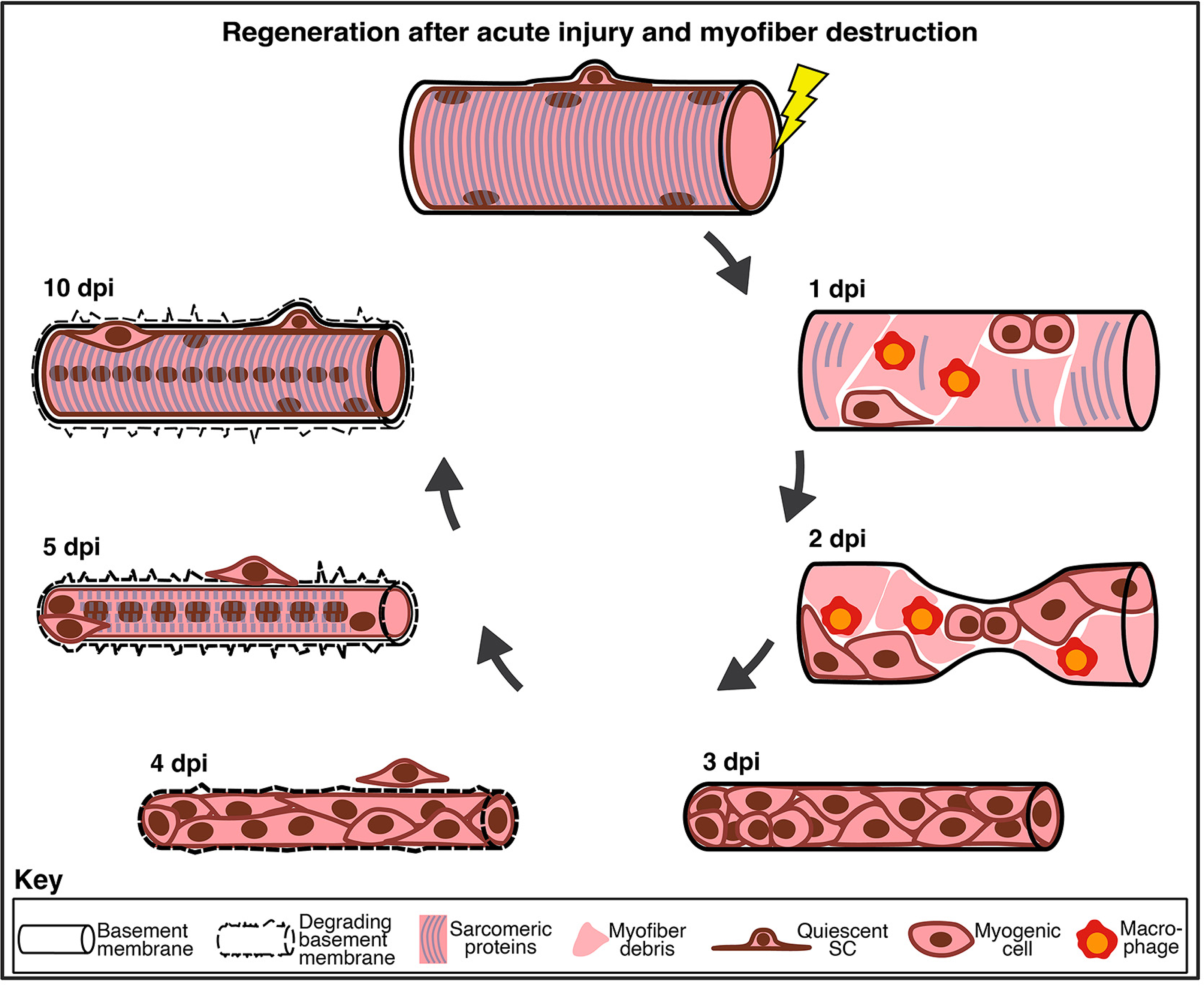
New model of cellular dynamics of regeneration. 1 DPI: Within residual BM, degradation of sarcolemma, macrophage entry and clearance of debris, and SC activation and proliferation. 2 DPI: Necking and shrinking of BM tube, within which macrophages clear debris and myogenic cells proliferate and are restricted to the periphery of BM by debris. 3 DPI: BM tube narrows and myogenic cells coalesce into columns. 4 DPI: BM tube narrowest and wave of density-dependent myocyte-myocyte fusion leads to myofibers with unaligned myonuclei. 5 DPI: Myofibers with myonuclei derived from myocyte-myocyte fusion positioned into centralized chains by forming sarcomeres and peripheral myonuclei derived from myocyte-myofiber fusion. New BM forming as old one degrades. 10 DPI: Myofibers continue to enlarge via occasional myocyte-myofiber fusion events and addition of peripheral myonuclei. Quiescent SCs are present in niche.

Activation and proliferation of SCs are the first events of the regeneration process. Our whole mount imaging found that prior to injury quiescent SCs are elongate, generally parallel to the myofiber, with delicate complex cellular projections — demonstrating both the sensitivity of our methods and confirming the findings of others ^41, 51–55^. When SCs enter Galert these cellular projections shorten and upon activation the projections are lost, findings consistent with recent work suggesting that the projections are critical for sensing myofiber damage and their retraction is one of the earliest events in SC activation ^54, 55^. Within 24 hours of activation, SCs enlarge, adopt a motile phenotype with lamellipodia, and begin proliferating.

SCs proliferate by a combination of symmetric and asymmetric divisions ^65^. There has been considerable debate as to when asymmetric cell division occurs and when SCs reacquire quiescence. Most previous research has used an ex-vivo system of SCs cultured on myofibers and based on the observation that the first division can be asymmetric, it has been proposed that early asymmetric division leads to early SC replenishment ^11, 59, 60, 63^. In addition, the inherent physical asymmetry of the SC niche ^5, 12, 44^ and observation of apical-basal divisions ex-vivo has led to the proposal that early apical-basal divisions generate asymmetric cell divisions that lead to SC replenishment ^11, 59, 60^. However, our in vivo characterization and quantification of SC division angles found only planar cell divisions and no apical-basal divisions 1-2 DPI and agrees with the results of Webster and colleagues ^52^. Together these data argue against a model whereby early apical-basal divisions are important for SC self-renewal. Two recent studies suggest that in vivo asymmetric cell division and SC self-renewal occur later in regeneration, generally at 5 DPI or later ^64, 65^. Consistent with these studies, we find that morphologically distinct quiescent SCs are not present until after 7 DPI, suggesting that the first priority for SCs is to regenerate myofibers and then only secondarily to re-establish a pool of quiescent SCs.

The main insights of our analysis are on the regeneration of myofibers and tissue architecture. Unexpectedly we uncovered that SCs and their myogenic derivatives regenerate myofibers via two distinct phases of fusion (Fig. 6). First, a massive wave of density-dependent myocyte-myocyte fusion regenerates myofibers 3.5-4.5 DPI. Given that the myogenic cells are sequestered within the BM tubes, it is likely that each nascent myofiber is derived from SCs associated with the damaged myofiber they are replacing. Notably, the myonuclei generated during this phase form the characteristic chains of centralized chains of myonuclei. Our experiments pulsing mice with EdU 4 hours prior to harvest demonstrate that after this primary phase of fusion, subsequently fusing myocytes never contribute to these centralized nuclei. Complementary studies with continuous BrdU ^34^ or EdU ^64^ labeling during multiple time windows during regeneration confirm that only myogenic cells labeled during 0-4 DPI contribute to centralized nuclei.

After this rapid primary wave of fusion, a second phase of fusion extends roughly from 5 to 10 DPI. This phase is density-independent and dominated by myocyte-myofiber fusion. These fusion events give rise to the peripheral myonuclei of regenerated myofibers, as found by our experiments labeling cells via pulses of TAM and/or EdU prior to harvest and also confirmed by the EdU and BrdU labeling experiments of ^34, 64^. Fusion of myocytes to myofibers is important for increasing the number of myonuclei and increasing the diameter of the regenerating myofibers. The appearance of myogenic cells outside the BM tubes beginning at 3.5 DPI suggests that some of these myocytes may derive from SCs originating from neighboring myofibers. While grafted SCs can migrate into adjacent myofibers ^75, 76^, it is currently unclear whether endogenous SCs only regenerate their home myofiber or whether they can contribute to regeneration of other myofibers (see discussion in ^13, 14^) and awaits future intravital imaging experiments.

Thus, myofibers are regenerated by two temporally and cellularly distinct mechanisms of fusion. Rapid primary myocyte-myocyte fusion quickly restores myofiber structure and function, while secondary myocyte-myofiber fusion optimizes regenerated myofibers. These two phases are likely to be governed by different molecular mechanisms as recent studies indicate that myocyte-myocyte and myocyte-myofiber fusion are differentially regulated ^77, 78^. Another important molecular mechanism question concerns the cessation of regeneration. We found that contribution of myocytes to myofibers is largely complete by 10 DPI, but what signals put a brake on fusion is unknown.

Our research also elucidates the etiology of the organization of myonuclei of regenerated myofibers. The distinctive centralized chains of myonuclei arise just after the wave of density-dependent fusion. Surprisingly, these chains do not result from successive planar cell division events, as had been predicted by ^34^, and they are also independent of innervation. Instead, they arise after fusion and are strongly correlated with myofibril assembly, suggesting that positioning of myonuclei is regulated by myofibrillogenesis. Previous studies have proposed a link between myofibril formation and myonuclei location ^67, 79, 80^, but in vitro studies have found that forming myofibrils drive nuclei to the myofiber periphery ^67^. However, in myofibers regenerated in vivo after a catastrophic injury, the initial myonuclei remain in centralized chains; we hypothesize that the rapidity of myocyte-myocyte fusion, nearly synchronous formation of myofibrils, and the enclosing BM traps and positions myonuclei into centralized chains.

The fate and persistence of centralized chains of myonuclei in regenerated myofibers has been considerably debated ^33, 36, 37^. Here we show that these chains remain for at least 10 months after injury, suggesting that they likely persist indefinitely and indelibly mark regenerated myofibers. The peripheral myonuclei that are commonly found in regenerated myofibers are not the result of peripheral movement of originally centrally located nuclei, but instead are derived from the later myocyte-myofiber fusion events. Thus, the common interpretation, particularly in the examination of diseased muscle, that centralized myonuclei mark actively regenerating myofibers needs to be revised ^37, 81^. An important consequence of this finding is that all myofibers regenerated after catastrophic myofiber damage will permanently have centralized myonuclei and may have compromised mechanical output. Intuitively, centralized myonuclei would seem to disrupt the function of sarcomeric proteins ^37^ and regenerated muscle overall has been demonstrated to be biomechanically stiffer than uninjured muscle ^82^, but whether individual regenerated myofibers with centralized myonuclei have an altered functional output has not been formally tested and awaits future experiments.

Our work highlights the importance of the residual BM for multiple aspects of muscle regeneration. During early stages of regeneration, the BM acts as a selective cell filter; BM tubes retain SCs and their myogenic derivatives, while allowing free penetration of infiltrating macrophages. Such a function had been suggested by older electron micrograph studies ^14, 19, 83^, but could only be demonstrated with our genetic labeling and whole mount imaging. The subsequent constriction of the BM (also observed in isolated regenerating human myofibers by ^79^) promotes the rapid reestablishment and proper orientation of myofibers. As revealed by electron micrography studies ^16, 17, 79^, the residual BM is gradually degraded and replaced by a new BM nested inside the old one, likely by 7 DPI (Fig. 6).

One of the key functions of macrophages during recovery from sterile muscle damage is the clearance of cellular debris (efferocytosis) ^70^. Efferocytosis is necessary to remove the remnants of damaged myofibers, allowing space for regeneration of new myofibers and also to induce the shift of macrophages from a pro-inflammatory to a pro-recovery phenotype ^70^. We found that debris clearance by pro-inflammatory macrophages is essential for the contraction of the residual BM, creation of space for proliferation of SCs and myoblasts, and unobstructed fusion of myocytes into linear myofibers. Depletion of macrophages and persistence of debris led to the formation of split and branched myofibers. Branched myofibers are characteristic of dystrophic muscle ^23, 28^, arise during regeneration ^28^, and are functionally deficient and more prone to damage compared with unbranched myofibers ^84, 85^. Several mechanisms have been proposed to cause branched myofibers, including aberrant fusion of myocytes ^18, 23, 31, 86^, defects in myogenic cell migration ^87^, or fusion of two myofibers ^88^. Here we show that macrophage-mediated removal of debris is essential for generating linear myofibers; without macrophages, cellular debris physically impedes fusion, resulting in branched myofibers. Thus, we find a new link between macrophage function and formation of pathogenic, branched myofibers.

Finally, our research elucidates similarities and differences between limb myogenesis during development and during regeneration. During both processes, myogenesis occurs in phases. The first phase (embryonic myogenesis or density-dependent myocyte-myocyte fusion in regeneration) establishes nascent myofibers, while the second phase (fetal myogenesis or myocyte-myofiber fusion in regeneration) enlarges these myofibers. The second phase is largely similar between development and regeneration; myogenesis proceeds over many days via myocyte-myofiber fusion resulting in peripheral myonuclei in myofibers surrounded by a BM. However, the first phase is quite different. In regeneration this phase happens through rapid density-dependent myocyte-myocyte fusion within a BM tube and results in chains of myonuclei that are locked into a central location within the myofibers. In contrast, embryonic myogenesis takes place in the absence of a laminin+ BM, extends over many days, is unlikely to be density-dependent, and results in centrally-located myonuclei which migrate to the myofiber periphery. These findings indicate that the molecular and cellular regulation of fusion ^78^ and myonuclear positioning ^37^ likely differ between development and regeneration.

In summary, our wholemount in vivo analysis of skeletal muscle after an acute, myofiber-destroying injury has provided significant new insights into the three-dimensional processes necessary to recover muscle structure (summarized in Fig. 6). Two distinct phases of fusion enable efficient restoration of fully functional myofibers, but indelibly mark regenerated myofibers with a distinct myonuclear architecture. Critical to this regenerative process is the residual BM; the BM spatially constrains SCs to promote rapid density-dependent myocyte fusion, orients and aligns myofibers, and (with collaboration of macrophages) ensures formation of linear unbranched myofibers. As centrally-nucleated and branched myofibers are characteristic of dystrophic muscle ^23, 28^, future whole mount analyses of mouse models of dystrophy will be important to test whether the normal regenerative processes are disrupted or amplified in disease. In most forms of experimentally-induced injury (e.g. BaCl_2_ and myotoxins) or naturally-occurring injury (e.g. muscle strain) the BM remains intact and the injury site remains sterile ^14, 15, 71^. However, in volumetric muscle loss the BM is removed in addition to the muscle ^89, 90^ and in pathogenic injuries (e.g. due to infection by viruses tropic to muscle) the macrophage and overall inflammatory response is altered ^71^. Future research that uses three-dimensional imaging to examine these forms of injury will provide insights into recovery after alternative forms of damage and further elucidate the general principles guiding regeneration of skeletal muscle, a tissue with a complex tissue architecture.

## Supporting information

CollinsVideoS1

CollinsVideoS2

CollinsVideoS3

CollinsVideoS4

CollinsVideoS5

## ACKNOWLEDGEMENTS

We thank Nathan Burns, Camila Goclowski, and Robert Krauss for reading drafts of the manuscript, and Y. Wan for help with Fluorender analysis. FACS was performed by the University of Utah Flow Cytometry core with assistance of James Marvin and confocal imaging was performed at the Cell Imaging Core with assistance of Xiang Wang. This work was supported by R01 HD104317, Pew Innovation Fund, Benning Endowed Chair to GK. MS was supported by NIH/NIAD T32 training grant AI055434.

## AUTHOR CONTRIBUTIONS

BBC, JS, and GK designed the experiments; BBC, JS, MS, RVM conducted experiments; BBC, JS, RVM, and GK analyzed the data; MV provided technical assistance; BBC and GK, with help from JS, wrote the paper.

## STAR METHODS

### RESOURCE AVAILABILITY

#### Lead contact

Further information and requests for resources and reagents should be directed to and will by fulfilled by the lead contact Gabrielle Kardon.

#### Materials availability

This study did not generate new unique reagents.

#### Data and code availability

All data reported in this paper will be shared by the lead contact upon request.

All original code and related data will be shared by the lead contact upon request.

Any additional information required to reanalyze the data reported in this paper is available from the lead contact upon request.

### EXPERIMENTAL MODEL AND SUBJECT DETAILS

#### Mice

Male and female 3-6 month *Pax7^CreERT^*^2 7^ and *Rosa^mTmG^* ^42^, back-crossed on to *C57/BL6J* background, as well as wild-type *C57/BL6J* mice (purchased from Jackson Laboratory) were used for all experiments in accordance protocols approved by the Institutional Animal Care and Use Committee at the University of Utah.

### METHOD DETAILS

#### Tamoxifen, EdU, and Encapsome/Clodrosome Injections

Tamoxifen (TAM, Cayman Chemical, 13258) doses of 2 mg were delivered by intraperitoneal injection. For EdU experiments 10μg/g body mass of EdU in PBS (ThermoFisher Scientific, A10044) was delivered by intraperitoneal injection. Macrophages were depleted by intraperitoneal injection of 150μl clodrosome (clodronate encapsulated liposome solution, Encapsula NanoSciences, CLD-8901) and controls were injected with encapsome (liposome solution without clodronate).

#### Muscle Injury

Barium chloride (25 μl 1.2% in sterile demineralized water) was injected into the TA or EDL via a Hamilton syringe, as in ^7^.

#### Common Peroneal Nerve Denervation

The common peroneal nerve was transected on the left hindlimb ^91–93^. A small skin incision was made in the direction from spine to thigh, centering over sciatic notch, and extending along the femur perpendicular to the course of the common peroneal nerve. Superficial fascia was incised to expose the hamstring muscles and a small incision was made through the hamstrings to expose the common peroneal nerve where it proximally intersects with the tendon of the lateral head of the gastrocnemius tendon. A 6-0 sterile silk suture was used to tie a knot at the proximal end and another at the distal end of the exposed nerve. The nerve was cut below the knot closest to the knee and the rest of the nerve was retracted by suturing to the bicep femoris. The hamstrings and skin incisions were closed with 6-0 sterile silk sutures.

#### FACS Analysis

Isolation of mononuclear myogenic cells and macrophages from the TA was performed as described previously ^94^. TAs from *Pax7^CreERT^*^2^*^/+^;Rosa^mTmG/+^* mice were dissected, minced and digested for 1 h at 37°C in 100 μl of 5mg/ml liberase (Sigma Aldrich, 5401127001) and 25 μl of 10U/µl DNAseI (Sigma Aldrich, 4716728001) in 3 ml Ham’s F12 media (ThermoFisher Scientific, 11765054). Samples were passed through 70um and 40um filters, spun at 1800 rpm for 10 min, supernatant aspirated, and pellet resuspended in SC growth media: 15% horse serum (Gibco, 16050-122), 1:1000 50mg/ml gentamicin (ThermoFisher Scientific, 15750060) in F12 media. Myogenic mononuclear cells were isolated via GFP. If needed, cells were incubated with Fc receptor CD16/CD32 (eBioscience, 14-0161-85) then stained with an antibody mixture of eFluor450 rat anti-mouse CD31(clone 390), PerCp-Cy5.5 rat anti-mouse CD45 (clone 30-F11), PeCy7 rat anti-mouse Sca-1 (clone D7), APC-eFluour780 rat anti-mouse F4/80 (clone BM8) and APC rat anti-mouse CD11b (clone M1/70) from eBiosciences, and Itga7 647 (clone R2F2) from AbLab (see Key Resources Table). Samples were incubated with antibodies on ice for 60 min, washed, and resuspended with SC growth media + DAPI for FACS analysis on the FACSAria III (BD Biosciences). Total mononuclear GFP+ myogenic cells (DAPI, TOM-negative, GFP+ or CD31-,CD45-, TOM-, GFP+) (Figure S1) and total macrophages (CD31-, CD45+, F4/80+, CD11b+) were calculated from entirely drained TA samples (Fig. S1). GFP+ cell diameter was measured using the flow cytometry size calibration kit from ThermoFisher Scientific.

#### Muscle Clearing for Whole-mount Imaging of Endogenous Fluorescence

The following protocol was adapted from ^41^. Mice were anesthetized with an overdose of isoflurane and an intracardiac perfusion and fixation was performed with PBS followed by 2% PFA. Harvested EDL samples were post-fixed in 2% PFA at 4°C 4 hours (H) - overnight. Fixed samples were washed 3 x 30 min in PBSTT: PBS with 0.1% Triton X-100 (Sigma Aldrich, T8787) and 0.1% Tween-20 (Sigma Aldrich, P7949). Samples were incubated in 4% acrylamide solution with 0.25% of thermal initiator VA-044 (Wako Chemical, 011-19365) for 4H rocking at 4°C. Samples were polymerized by incubated in fresh, degassed acrylamide solution for 3-4 H rotating at 37°C. The polymerized samples were washed 3 x 30 min with PBSTT at 37°C. Samples were then incubated in clearing solution (pH 9.3) containing 10% N,N,N’,N’-tetrakis ethylenadiamine (Sigma-Aldrich, H2383), 10% Urea (Bio-Rad, 1610731), 5% Triton X-100, 5% sodium deoxycholate (Sigma Aldrich, D6750), and 20mM boric acid (Sigma Aldrich, B6768) rotating overnight at 37°C. Prior to mounting samples for confocal imaging, the refractive index of the tissue was matched using 88% Histodenz in PBSTT (Sigma Aldrich D2158) with 0.01% w/v sodium azide (Sigma Aldrich S2002) overnight at RT. Samples were submerged in 88% Histodenz solution in a well slide for imaging. Samples were protected from light during the entire protocol.

#### Whole-mount Immunostaining

##### GFP, TOM, Laminin, Myosin, and EdU

After the modified muscle tissue clearing protocol from ^41^, samples were washed extensively in PBS at RT and then incubated 1H at RT in blocking solution containing 5% goat serum (ThermoFisher Scientific, 16210-072), 20% DMSO (Fisher Scientific, D159-4) in PBS with anti-mouse Fab fragments (Jackson ImmunoResearch, 115-007-003) added if needed. Then samples were incubated with primary antibodies diluted in 5% goat serum and 20% DMSO in PBS for 2 days at RT; primary antibodies included GFP (1:500; Aves Labs, GFP-1010), and dsRed (1:200; Takara, 632496), laminin (1:100; Sigma Aldrich, L9393), and Myosin Heavy Chain (1:000 Sigma Alrich M4276). Samples were washed extensively with PBS and then incubated in secondary antibody diluted in 5% goat serum and 20% DMSO in PBS for 3 days at RT in the dark; secondary antibodies were conjugated with Alexa 488, Alexa 594, or Alexa 647 (Jackson ImmunoResearch; see Key Resources Table). Samples were washed extensively in PBS and stored at 4°C in the dark until imaging. Samples were washed 3 x 30 min with PBSTT prior to incubation in Histodenz solution.

Samples labeled with EdU were additionally blocked in 3% BSA in PBS for 15 min at RT and then underwent the Click-it reaction at 37°C according to manufacturer’s protocol. Click-it reaction was quenched with 15 min of 3% BSA in PBS followed by extensive PBSTT washes at RT.

##### CD68

The following protocol was adapted from ^95^. Mice were anesthetized with an overdose of isoflurane and an intracardiac perfusion and fixation was performed with PBS followed by 4% PFA. Harvested EDL samples were post-fixed in 4% PFA at 4°C 4H - overnight. Fixed samples were washed extensively in PBS and dehydrated with a series of 20%, 40%, 60%, 80% methanol in H_2_O with 0.1% Triton X-100 and 0.3 M glycine (B1N buffer, pH 7) and 100% methanol washes for 30 min each at 4°C shaking. Samples were then delipidated in 100% dichloromethane (DCM, Sigma Aldrich, 270997) for 3 x 30 min at 4°C with shaking. Samples were washed with 100% methanol three times and then rehydrated in a reversed methanol/B1N series, 80% 60%, 40%, 20% for 30 min each at 4°C. Samples were then washed in B1N 2 x 30 min at RT and washed overnight in B1N at RT. Samples were permeabilized in PBS with 0.1% Triton X-100, 0.05% Tween20, and 2μg/ml heparin (PTwH) for 2 x 1H at RT. Samples were incubated in primary antibody in PTwH for 3 days at RT. The primary antibodies used with this protocol were GFP (1:500; Aves Labs, GFP-1010) and CD68 (1:100; Bio-Rad, MCA1957). Samples were washed extensively with PTwH and then incubated in secondary antibody diluted in PTwH for 3 days at RT in the dark; secondary antibodies were conjugated with Alexa 488 and Alexa 647 (Jackson ImmunoResearch; see Key Resources Table). Samples were washed extensively in PTwH prior to tissue clearing. To clear the tissue, samples were dehydrated in 25%, 50%, 75%, 100%, 100%, methanol/H_2_O series for 30 min each at RT. Following dehydration, samples were washed with 100% DCM for 30 min three times, followed by overnight clearing in dibenzyl ether (DBE; Sigma Aldrich, 10814) at RT. Samples were mounted submerged in 100% DBE in a welled slide for imaging

#### Whole-mount Imaging and 3D Rendering

Samples were imaged on a Leica SP8 confocal microscope with PL APO 20x/0.75 and 63x/1.40 oil immersion objectives were used with Z optical sections of 3 μm and 1 μm, respectively. A linear increase in laser intensity was used for deeper tissue imaging.

Optical stacks of images were rendered in depth mode with Fluorender ^46^. To highlight objects (GFP+ cells, basement membrane, branched fibers), objects were selected in Fluorender using the paint brush tool on individual Z optical sections and objects were extracted, rendered and pseudo-colored.

#### Quantification of Myogenic Cell and BM Morphology

##### Myogenic Cell Volume

In Fluorender GFP+ cells were selected using the paint brush tool on individual Z optical sections and extracted and the component analyzer tool was used to measure cell volume of the extracted cells.

##### SC Projection Number and length

In Fluorender individual GFP+cells were extracted and the polyline ruler tool was used measure projection length. Projections were measured from the nucleus to the tip of the longest branch; 3 μm was the minimum length required to be considered a projection. Because many projections split and have several terminal ends, branches longer than 3 μm were measured from the junction with the main projection to tip of the branch. The length of all the branches on one projection were summed to give the total projection length.

##### Myofiber Diameter

In Fluorender, images of GFP+ regenerated myofibers were rotated to view myofibers in cross-section. Then images of cross-sections at multiple points along the length of the myofibers were obtained. These images were imported into Image J and myofiber diameter was measured using the line tool at each y-axis cross-section and averaged for each myofiber analyzed.

##### BM Diameter

Using Fluorender, the diameter of 3D rendered images of laminin+ BM tubes were measured at multiple points along the length of the tubes using the ruler tool. Then the average diameter was calculated for each BM tube measured. At 2dpi, the widest and most constricted dimensions of BM tubes was measured with the ruler tool.

##### Branched Fibers

To determine the percentage of branched regeneration fibers in each muscle sample, Z-stacks of the composite rendered images were scanned for branched regenerated GFP+ myofibers. If there were multiple branches on a single myofiber, this was still counted as one branched myofiber. Because there was variability in the number of regenerated myofibers between samples, myofiber branching was expressed as a percentage of total regenerated GFP+ myofibers. The total number of regenerated myofibers was counted on images rotated to visualize myofibers in cross-section.

### QUANTIFICATION AND STATISTICAL ANALYSIS

Statistical analyses were performed using Prism 9 (Graphpad, La Jolla, CA). For comparison between two groups, unpaired t tests were used. For comparison between multiple groups, one-way ANOVA with multiple comparisons Holm-Sidak post-hoc analysis was used. An alpha level of 0.05 was used for all analyses. All values are displayed as means ± SEM. Asterisk (*), (**), (***), (****) indicate statistically significant differences between the compared values (p < 0.05, p < 0.01, p < 0.001 and p < 0.0001, respectively).

## SUPPLEMENTAL FIGURE AND VIDEO LEGENDS

**Figure S1.**
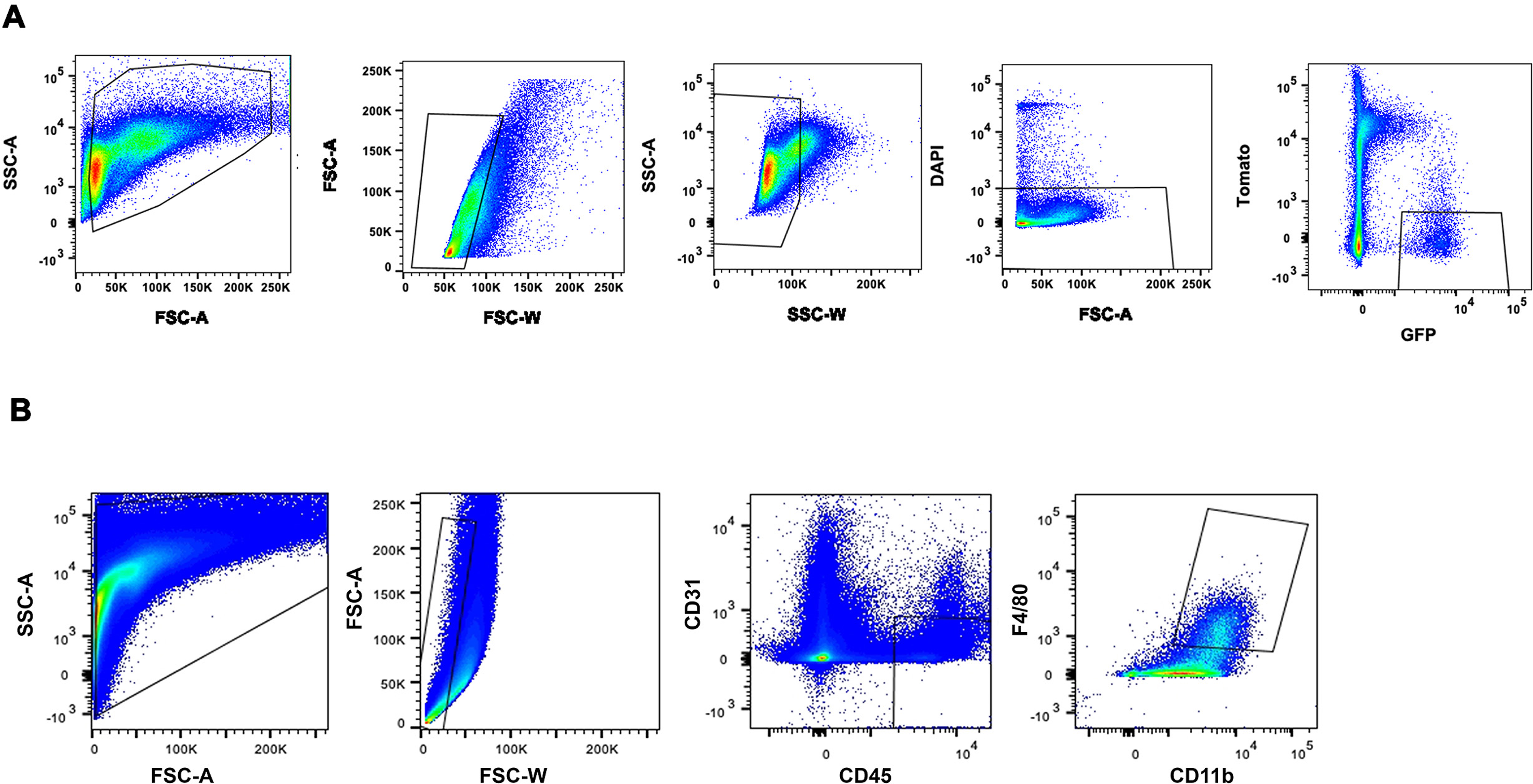
Gating strategy for GFP+ mononuclear cells and macrophages. **(A)** TA from adult *Pax7^creER/+^;Rosa^mTmG/+^*mice were enzymatically digested and gated based on forward/side scatter (plots 1-3) and live cells (DAPI, plot 4) and then selected for Tomato- and GFP+ (plot 5). **(B)** Muscles from *C57BL/6J* mice were enzymatically digested and gated based on forward/side scatter (plots 1-2), CD31- and CD45+ cells (plot 3), and double positive for F4/80 and CD11b (plot 4).

**Figure S2.**
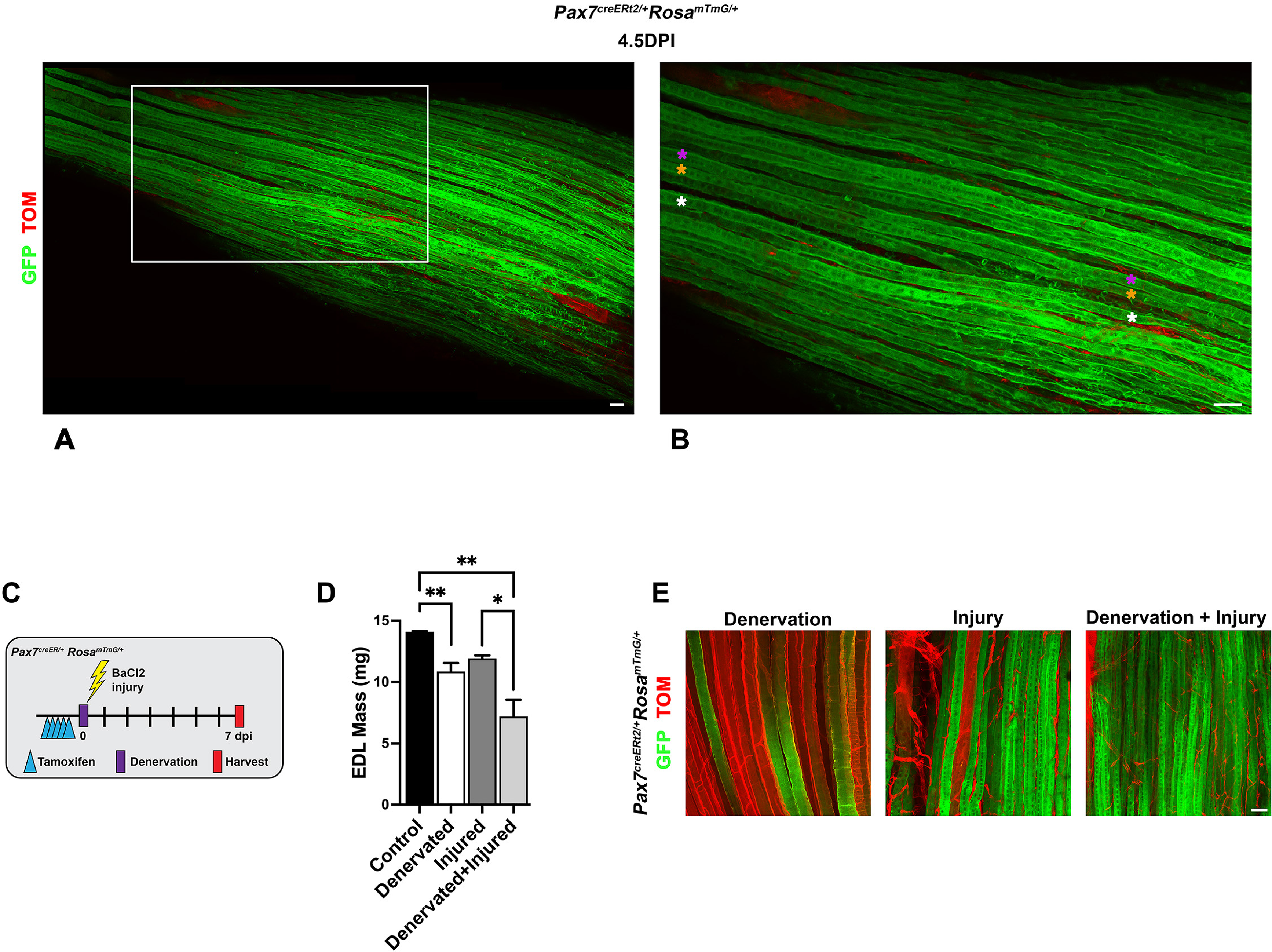
Formation of centralized chains of myonuclei occurs after fusion and independent of innervation. **(A-B)** Representative tile scan of EDL at 4.5 DPI (A) with zoomed region in (B). Pink, yellow, and white asterisks indicate single myofibers with chains of myonuclei on left halves and unaligned nuclei on right halves. **(C-E)** After denervation of common peroneal nerve and/or BaCl_2_ injury (experimental scheme, C), muscle mass decreases with denervation (D) but centralized chains still are present 14 DPI (E). Histogram values are mean ± SEM; * p < 0.05, ** p < 0.01. (n = 3 mice/experiment). Scale bar on all images = 50um.

**Video S1. Quiescent, Galert, and activated SCs are distinct in size and morphology.** Representative wholemount reconstructions of uninjured, Galert (2.5 DPI contralateral uninjured EDL),1 DPI, and 1.5 DPI EDLs. Scale bar on all images = 50um.

**Video S2. Regenerated myofibers form via a wave of density-dependent fusion 3.5-4.5 DPI followed by alignment of myonuclei by 5 DPI.** Representative wholemount reconstructions of 2, 2.5, 3, 3.5, 4, 4.5, and 5 DPI EDLs. Scale bar on all images = 50um.

**Video S3. Myofibers enlarge by contribution of myogenic cells to peripheral nuclei.** Representative wholemount reconstructions of 7, 10, 14, and 21 DPI EDLs. Scale bar on all images = 50um.

**Video S4. Development of myofibers is distinct from regeneration; myogenesis begins in the absence of continuous BMs and initially centralized myonuclei move peripherally.** Representative whole reconstructions of E12.5, E14.5, and P0 TAs. Scale bar on all images = 50um.

**Video S5. Macrophages are essential to clear debris, enabling density-dependent myogenic cell fusion and preventing formation of branched myofibers.** Representative whole mount reconstructions of 2.5 DPI showing myogenic cells and macrophages within BM tube at 2.5 DPI . Also representative whole mount reconstructions 4 and 7 DPI showing that macrophage depletion leads to poorly fused and branched myofibers. Scale bar on all images = 50um.

